## supplementary for "Aging drives cerebrovascular network remodeling and functional changes in the mouse brain"

### Supplementary Figure 1

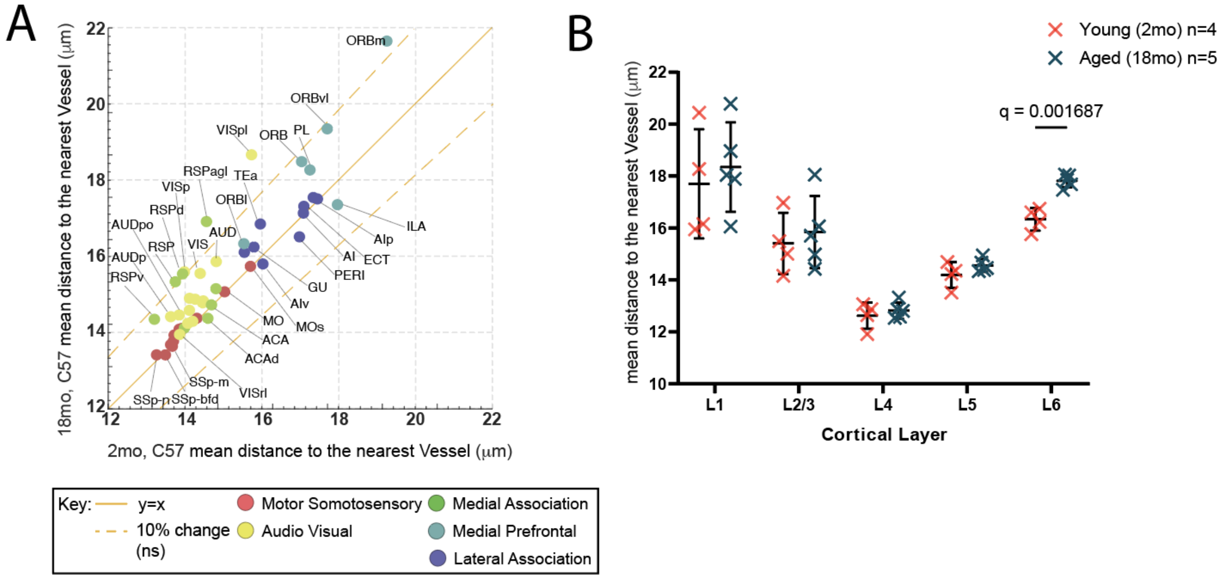


**Nearest neighbor distance (NND) to vessels**. (**A**) Scatter plots of averaged NND (μm) in isocortical areas. No areas show significant differences. (**B**) NND showed significant reductions only in layer 6. Brain region abbreviations can be found in Supplementary Data 1.

### Supplementary Figure 2

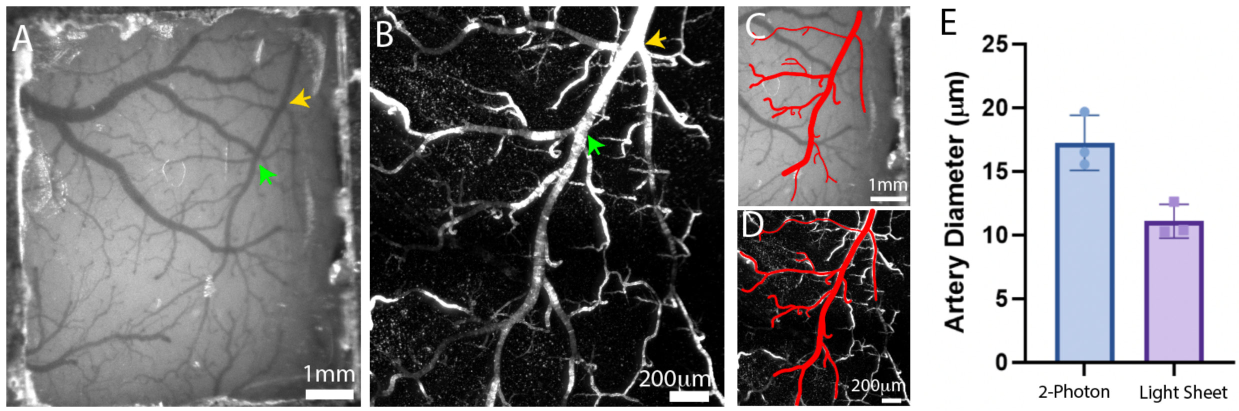


**Intact vascular geometry after tissue clearing and immunolabeling**. (**A-B**) In vivo two-photon imaging (**A**) and light sheet imaging (**B**) of the matched area from the same animal. Yellow and green arrows indicate matching vessels in the two-photon and light sheet imaging. (**C-D**) A selection of an artery (red) as an example in the two-photon imaging (**C**) and its overlay with scaling in the light sheet imaging (**D**). Note the near complete overlap. (**E**) Artery diameter measurement from the same artery shows ~36% decrease in the light sheet imaging after the tissue clearing.

### Supplementary Figure 3

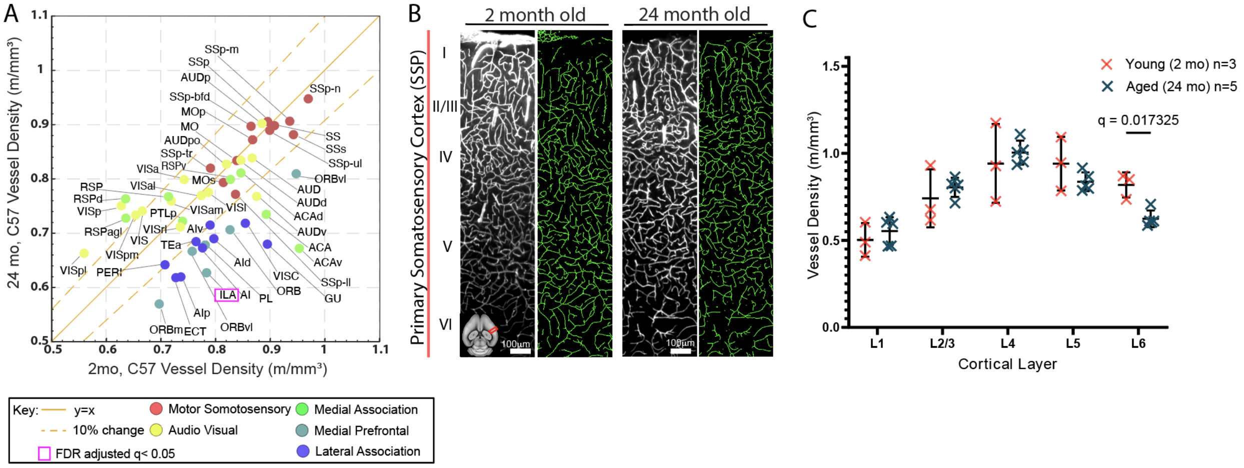


**Selective reduction of vascular length density in the late aging mice**. (**A**) Scatter plots of vascular length density (m/mm$^3$) in isocortical areas from 2-month-old and 24-month-old mice by light sheet imaging. Only the infralimic cortex shows a significant difference. (**B**) Examples of light sheet imaging with lectin pan-vascular staining (left) and tracing (right, green) from 2-month-old and 24-month-old mice. (**C**) Vascular length density across cortical layers showed that only layer 6 showed significant reduction in the aged brain.

### Supplementary Figure 4

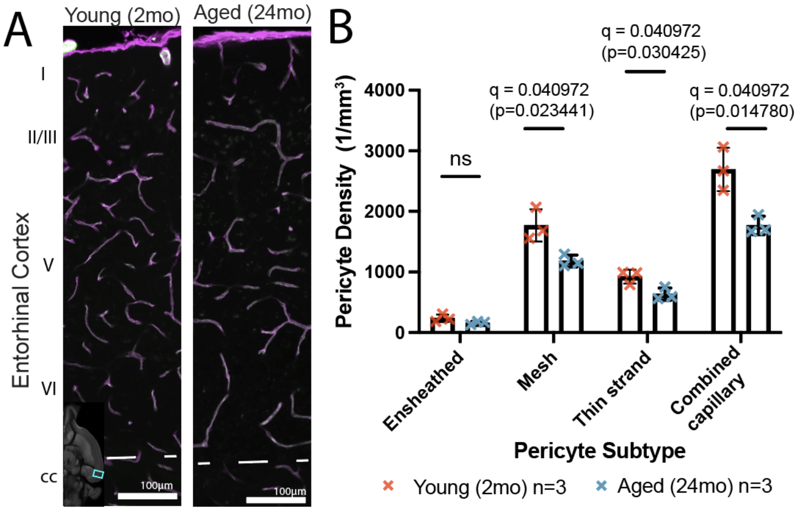


**Significant reduction of pericytes in the entorhinal cortex.** (**A-B**) The entorhinal cortex (**A**) showed a significant reduction of capillary pericytes (**B**).

### Supplementary Figure 5

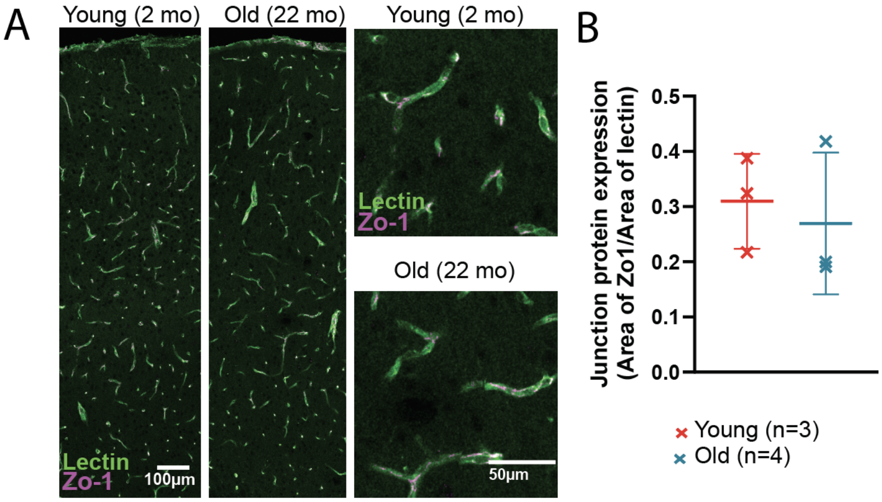


**Lack of significant changes in the Zo-1 expression in the aged brain.** (**A**) Zo-1 and Lectin staining in 2-month-old and 22-month-old brains. (**B**) No significant difference between the two age groups.

### Supplementary Figure 6

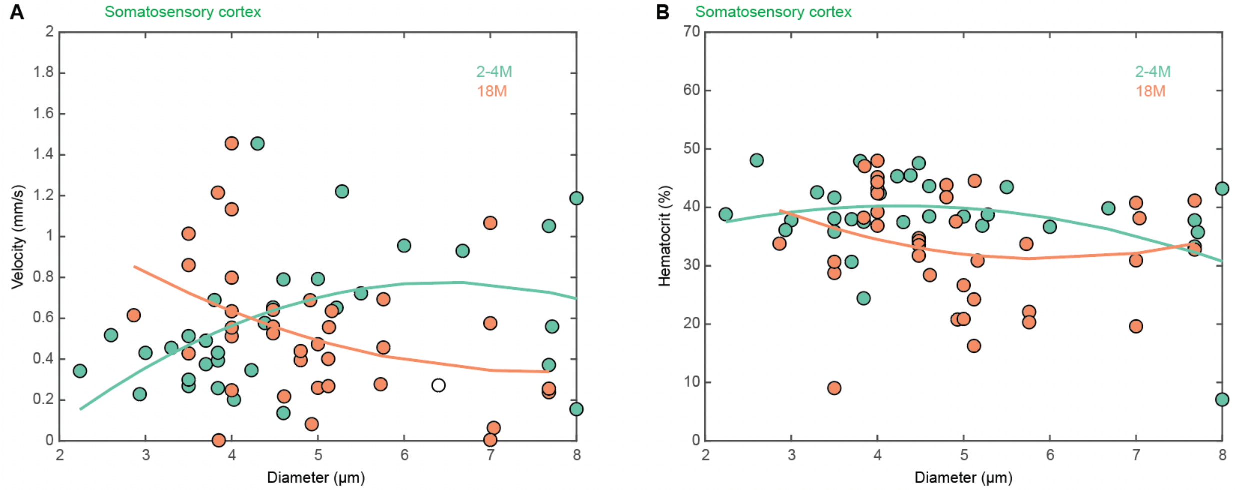


**Red blood cell velocity and hematocrit in the capillary network**. (**A**) Scatter plots showing basal RBC velocity as a function of capillary lumen diameter. Solid lines indicate a least mean square fit for all the data points in each group. (**B**) As in (**A**) but for instantaneous hematocrit.

**Supplementary Text 1. Red blood cells spacing in the capillary network does not change with aging.**

We quantified the spacing of RBC and compared the occurrence of "stall" events during different aging groups. Approximately 207 minutes data from 68 capillaries in 13 mice were analyzed. In young (2-4 month old) mice, for all the RBC intervals during long resting periods (approximately 104 minutes data from 32 capillaries in 8 mice; 29.4 ± 18.6 ms, median ± interquartile range; 95% confidence interval: [16.6 ms, 122.2 ms]), only approximately 0.02% RBC intervals are stall events (1.41 ± 1.37 second, median ± interquartile range; 95% confidence interval: [1.01 second, 6.56 seconds]). In 18-month old mice, for all the RBC intervals during long resting periods (approximately 103 minutes data from 36 capillaries in 5 mice; 31.4 ± 23.7 ms, median ± interquartile range; 95% confidence interval: [16 ms, 107.3 ms]), approximately 0.02% RBC intervals are stall events (1.77 ± 1.71 second, median ± interquartile range; 95% confidence interval: [1.00 second, 12.6 seconds]). These results suggest that there are no significant changes in stall occurrence in healthy aging, although mouse models of Alzhemier's disease show significant more occurrence of stall events.
